# Divergent polo boxes in KKT2 and KKT3 initiate the kinetochore assembly cascade in *Trypanosoma brucei*

**DOI:** 10.1101/2022.06.02.494600

**Authors:** Midori Ishii, Patryk Ludzia, Gabriele Marcianò, William Allen, Olga O. Nerusheva, Bungo Akiyoshi

## Abstract

Chromosome segregation requires assembly of the macromolecular kinetochore complex onto centromeric DNA. While most eukaryotes have canonical kinetochore proteins that are widely conserved among eukaryotes, evolutionarily divergent kinetoplastids have a unique set of kinetochore proteins. Little is known about the mechanism of kinetochore assembly in kinetoplastids. In this study, we have characterized two homologous kinetoplastid kinetochore proteins, KKT2 and KKT3, that constitutively localize at centromeres and promote the recruitment of other kinetochore proteins. KKT2 and KKT3 have three domains that are highly conserved among kinetoplastids: an N-terminal kinase domain of unknown function, the centromere localization domain in the middle, and the C-terminal domain that has weak similarity to polo boxes of Polo-like kinases. We show that the kinase activity of KKT2 is essential for accurate chromosome segregation, while that of KKT3 is dispensable for cell growth in the procyclic form of *Trypanosoma brucei*. Crystal structures of their divergent polo boxes reveal differences between KKT2 and KKT3. We also show that the divergent polo boxes of KKT3 are sufficient to recruit KKT2 in *T. brucei*. Furthermore, we identify the C-terminal domain of KKT1 as a direct interaction partner for the divergent polo boxes of KKT2, but not KKT3. KKT6 is found to interact with KKT1. These results show that KKT2 and KKT3 recruit downstream kinetochore proteins by direct protein-protein interactions using their divergent polo boxes.

## Introduction

The kinetochore is the macromolecular protein complex that drives chromosome segregation in eukaryotes (McIntosh, 2016). It assembles onto centromeric DNA and interacts with spindle microtubules during mitosis and meiosis. In most eukaryotes, the position of kinetochore assembly sites is marked by the presence of a centromere-specific histone H3 variant, CENP-A, which also plays important roles in the recruitment of other kinetochore proteins (Musacchio and Desai, 2017). CENP-A is found in diverse eukaryotes, suggesting that most eukaryotes use CENP-A to initiate the kinetochore assembly cascade (Drinnenberg and Akiyoshi, 2017; van Hooff et al., 2017). However, CENP-A is absent in some lineages such as kinetoplastids, holocentric insects, and early-diverging fungi, implying that there are alternative mechanisms for kinetochore assembly (Drinnenberg et al., 2014; Navarro-Mendoza et al., 2019; Ishii and Akiyoshi, 2022).

Kinetoplastids are flagellated eukaryotes defined by the presence of a unique organelle called the kinetoplast that contains a cluster of mitochondrial DNA (d’Avila-Levy et al., 2015). They are evolutionarily divergent from traditional model eukaryotes used in kinetochore research, such as yeasts, worms, flies and humans (Keeling and Burki, 2019). Interestingly, none of canonical kinetochore proteins including CENP-A has been found in the genome of kinetoplastids (Lowell and Cross, 2004; Berriman et al., 2005; van Hooff et al., 2017). Instead, experimental studies in *Trypanosoma brucei* have identified unique kinetochore proteins called KKT1–25 (Akiyoshi and Gull, 2014; Nerusheva et al., 2019) and KKIP1–12 (D’Archivio and Wickstead, 2017; Brusini et al., 2021), many of which are conserved in kinetoplastids (Butenko et al., 2020; Geoghegan et al., 2021). Dissecting the unique kinetoplastid kinetochores can provide insights into the evolution and fundamental requirements of chromosome segregation in eukaryotes. Based on architectural similarities between kinetoplastid kinetochores and synaptonemal complexes as well as sequence similarities between their components, we proposed that kinetoplastids might have repurposed meiotic synaptonemal complexes and homologous recombination machinery to assemble the unique kinetochore (Tromer et al., 2021).

There are a number of outstanding questions about kinetoplastid kinetochores. For example, it remains unknown how kinetoplastid kinetochores assemble on centromeres. In other eukaryotes, CENP-A-containing nucleosomes recruit components of the constitutive centromere-associated network (CCAN) to initiate the kinetochore assembly cascade. There are six proteins in *Trypanosoma brucei* that constitutively localize at kinetochores (KKT2, KKT3, KKT4, KKT20, KKT22, and KKT23), and these proteins may form the foundation of kinetoplastid kinetochores (Akiyoshi and Gull, 2014; Nerusheva et al., 2019). Growth defects have been observed upon depletion of KKT2, KKT3, KKT4, and KKT23 (Marcianò et al., 2021). KKT4 has microtubule-binding activity (Llauró et al., 2018; Ludzia et al., 2021), while KKT23 has a GCN5-related N-acetyltransferase domain of unknown function (Nerusheva et al., 2019). Importantly, kinetochore localization of KKT2 and KKT3 was not affected when KKT4 or KKT23 were depleted, while depletion of KKT2 and KKT3 affected the localization of other kinetochore proteins including KKT4 and KKT23 (Marcianò et al., 2021). Together with that finding that KKT2 and KKT3 have DNA-binding motifs (Akiyoshi and Gull, 2014), it is plausible that these constitutive kinetochore proteins form the base of kinetoplastid kinetochores and recruit other kinetochore proteins to initiate the assembly cascade.

KKT2 and KKT3 are homologous to each other and have several domains that are highly conserved among kinetoplastids, including an N-terminal kinase domain, the central domain, and divergent polo boxes (DPB) (Figure 1A) (Nerusheva and Akiyoshi, 2016). The kinase domain of KKT2 and KKT3 has been classified as unique among eukaryotic kinases (Parsons et al., 2005), and very little is known about the function or substrate of these kinase domains. The central domain of KKT2 and KKT3 localized at kinetochores when ectopically expressed in trypanosomes and was named the centromere localization domain (Marcianò et al., 2021). Polo boxes are protein-protein interaction domains found in Polo-like kinases (PLKs) that often must be phosphorylated to enable protein-protein interactions (Elia et al., 2003; Zitouni et al., 2014). Immunoprecipitation of ectopically expressed KKT2 DPB and KKT3 DPB from trypanosomes revealed co-purification of several kinetochore proteins, supporting a possibility that these divergent polo boxes are protein-protein interaction domains that contribute to kinetochore assembly in kinetoplastids (Marcianò et al., 2021). To date, direct interaction partners of the divergent polo boxes remain unknown.

**Figure 1.**
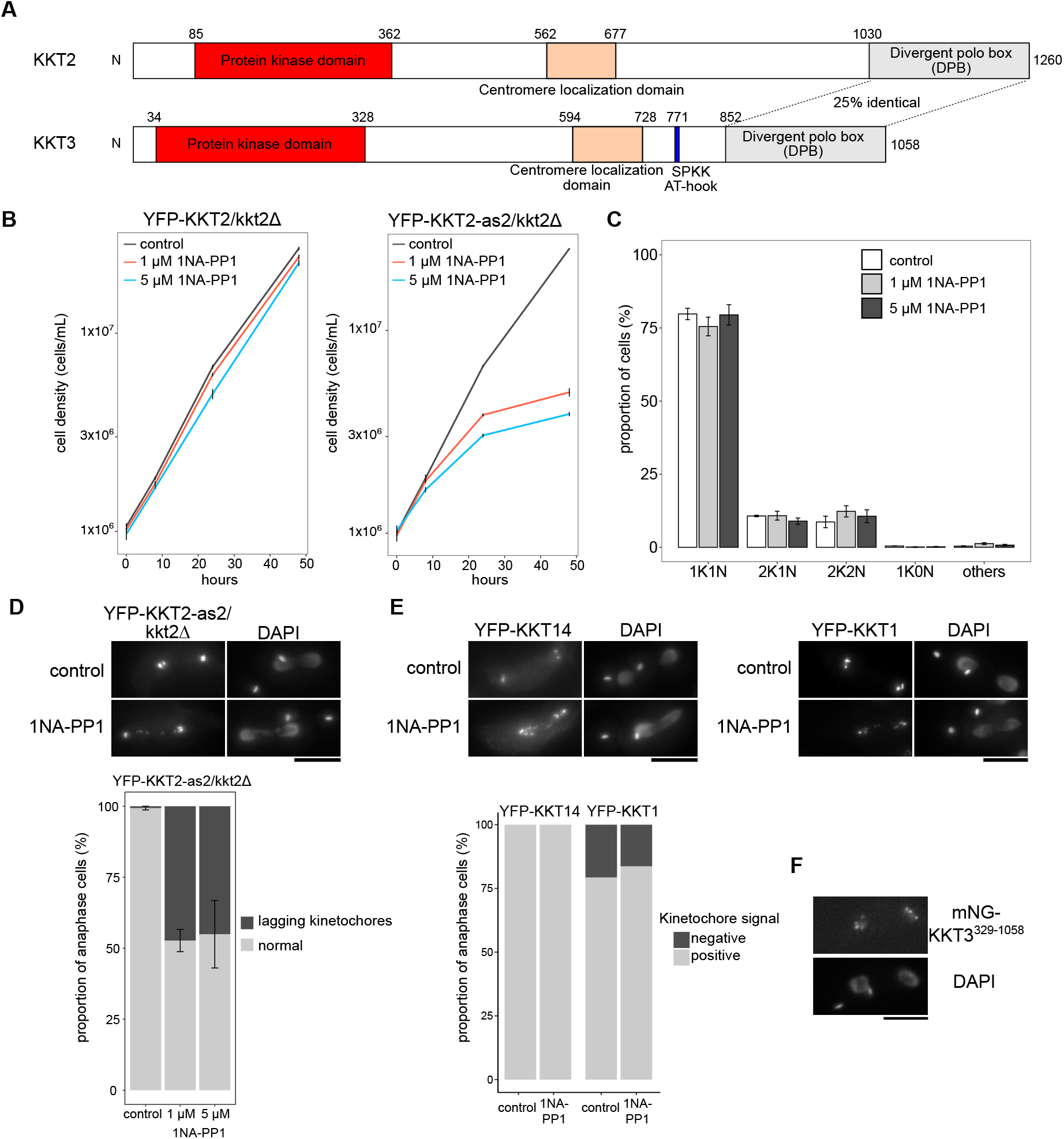
Kinase activity of KKT2 is essential for accurate chromosome segregation. (A) Schematics of *T. brucei* KKT2 and KKT3, highlighting the N-terminal unique kinase domain, the centromere localization domain in the middle, and C-terminal divergent polo boxes, as well as the SPKK and AT-hook DNA-binding motifs in KKT3. (B) Procyclic cells that have a wild-type KKT2 copy are not sensitive to 1NA-PP1 (left), while cells that have a KKT2-as2^M161A^ analogue-sensitive allele impairs cell growth upon addition of 1 µM or 5 µM 1NA-PP1 (right). Control is DMSO-treated cells. Error bars show SEM (n=3). Cell lines: BAP963 and BAP2269 (C) Cell cycle counts after 1NA-PP1 treatment for 8 hours, showing no significant difference in the cell cycle profile (p-value > 0.05). Error bars show SEM (n=3, >240 cells each). Cell line: BAP2269 (D) Inhibition of KKT2 kinase activity causes chromosome segregation defects. Images of lagging kinetochores (top) and quantification of lagging kinetochore (bottom) are shown. KKT2 analogue-sensitive cells were treated with 1 µM 1NA-PP1, 5 µM 1NA-PP1, or DMSO for 8 hours and fixed for microscopy. Cells treated with 1NA-PP1 have significantly more lagging kinetochores than untreated control (p-value < 0.0001, Fisher’s exact test for total numbers of cells from three independent experiments. p-value < 0.0001, Fisher’s exact test for count data [control vs 1 µM] and [control vs 5 µM]. Error bars show SEM (n=3, >42 anaphase cells each). Cell line: BAP2269 (E) Kinetochore localization of KKT14 and KKT1 is not affected by inhibition of KKT2 kinase activity. Top: KKT2 analogue-sensitive cells expressing YFP-KKT14 (left) or YFP-KKT1 (right) were treated with 5 µM 1NA-PP1 or DMSO for 8 hours and fixed for microscopy. Bottom: Quantification of cells that had kinetochore signals. Anaphase cells were analyzed (>18 cells each). Cell lines: BAP806 and BAP839 (F) Procyclic cells can survive without KKT3 kinase domain. Cells expressing mNeonGreen-KKT3^329–1058^ as the sole copy of KKT3 were fixed, showing normal kinetochore localization. Cell line: BAP1014. Bars = 5 µm.

Here we report characterization of the kinase domain and divergent polo boxes of KKT2 and KKT3. We show that the kinase activity of KKT2, but not KKT3, is essential for accurate chromosome segregation in the procyclic form of *Trypanosoma brucei*. Crystal structures of their divergent polo boxes confirm similarity to the polo boxes of PLK1 and reveal differences between KKT2 and KKT3. We also show that KKT3’s divergent polo boxes can recruit KKT2 in trypanosomes and that KKT2’s divergent polo boxes directly interacts with KKT1. These results suggest that KKT2 and KKT3 play key roles in the assembly of kinetoplastid kinetochores by recruiting downstream kinetochore proteins using their divergent polo boxes.

## Results

### KKT2 kinase activity is essential for chromosome segregation

A recent study showed that a kinase-dead allele of KKT2 was unable to support proliferation of the bloodstream form *T. brucei* (Saldivia et al., 2021). To examine the importance of the KKT2 kinase activity in the procyclic form, we created analog-sensitive KKT2 alleles by mutating a gatekeeper residue (M161) (Bishop et al., 2000). The cell line carrying KKT2-as2 (M161A) as the sole copy of KKT2 grew normally but had a growth defect upon addition of 1NA-PP1, an analog of the PP1 kinase inhibitor (Figure 1B). Judging from the number of kinetoplasts (K) and nuclei (N) in a cell (Robinson et al., 1995), there was no striking change in the cell cycle profile upon inhibition of the KKT2 kinase activity (Figure 1C). Kinetochore localization of KKT2 was not affected by inhibition of its kinase activity, but >40% of anaphase cells had lagging kinetochores after an 8-hour treatment (Figure 1D). These results showed that the kinase activity of KKT2 is essential for proper chromosome segregation in the procyclic form. We next examined the contribution of KKT2’s kinase activity for the localization of other kinetochore proteins. Based on our previous finding that KKT14 failed to localize at kinetochores upon RNAi-mediated depletion of KKT2 (Marcianò et al., 2021), we first monitored the localization of KKT14 but found that it still localized at kinetochores upon inhibition of the KKT2 kinase activity (Figure 1E). KKT1 localization was also unaffected (Figure 1E). These results show that the kinase activity of KKT2 is essential for accurate chromosome segregation but is dispensable for the localization of itself, KKT1, and KKT14.

### KKT3 kinase activity is dispensable for the proliferation of procyclic cells

Similarly to KKT2, KKT3 is essential for cell growth (Akiyoshi and Gull, 2014; Jones et al., 2014). To our surprise, cells carrying KKT3-as1 (M109G) as the sole copy of KKT3 did not have any growth defect even in the presence of 1NA-PP1, 1NM-PP1, or 3MB-PP1 (see Materials and Methods). We next created a kinase-dead allele of KKT3 by mutating lysine 63 (a residue conserved in active protein kinases in eukaryotes) and found that the cell line carrying KKT3^K63A^ as the sole copy of KKT3 grew normally without obvious defects (data not shown). We even obtained a cell line that entirely lacked the kinase domain of KKT3 (Figure 1F). These results show that the kinase activity of KKT3 is not essential for cell growth in procyclic cells. Given its high level of sequence conservation, however, it is possible that its kinase activity is essential under certain growth conditions or life stages.

### KKT2 phosphorylates KKT8, while KKT3 phosphorylates KKT12

We next aimed to identify substrates of KKT2 and KKT3 by in vitro kinase assays using recombinant proteins (Figure S1A). While screening several kinetochore proteins and histones, we found the KKT8 complex that consists of KKT8/9/11/12 proteins (Ishii and Akiyoshi, 2020) as an in vitro target, where KKT2 phosphorylated KKT8, while KKT3 phosphorylated KKT12 (Figure 2A). By mutating conserved serine/threonine residues, we identified S381 as the major site in KKT8 targeted by KKT2 (Figure 2B), while phosphorylation of KKT12 by KKT3 was significantly reduced when either T188 or S192 were mutated (T188A, T188S, and S192A) (Figure 2C). It is possible that S192 is the major phosphorylation site and that T188 is required for efficient phosphorylation of S192. An alternative possibility is that T188 is the major phosphorylation site and S192 is required for efficient phosphorylation of T188. These sites and surrounding residues are highly conserved among kinetoplastids (Figure 2D, 2E, Figure S2, and Figure S3). It is noteworthy that the −2 position (residue 379 in KKT8) is glycine (Figure 2D), which is not commonly found in the consensus phosphorylation motif of protein kinases (Hutti et al., 2004). Interestingly, our previous work showed that the BRCT domain of KKT4 binds the KKT8 peptide and that phosphorylation of S381 increased the affinity (Ludzia et al., 2021). These results raise a possibility that KKT2 may regulate the KKT4 BRCT domain (of unknown function) through phosphorylation of KKT8. To examine the importance of this phosphorylation event, we performed a rescue experiment in trypanosomes but found that the KKT8^S381A^ mutant did not affect cell growth (Figure 2F). Because inhibition of the KKT2 kinase activity causes growth defects (Figure 1B), this result means that KKT2 has additional targets to promote accurate chromosome segregation. More work is needed to identify such substrates.

**Figure 2.**
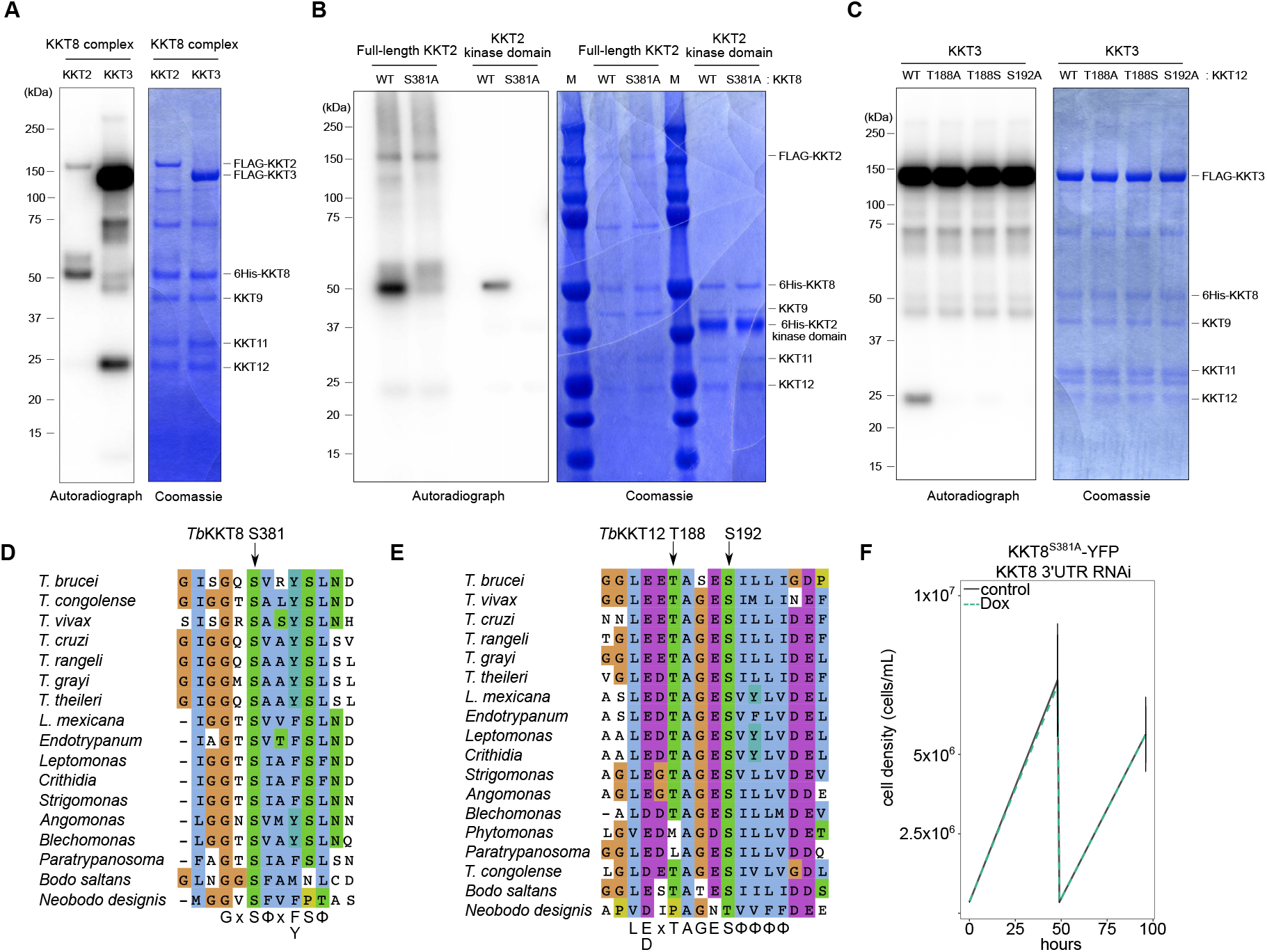
KKT2 phosphorylates KKT8, and KKT3 phosphorylates KKT12. (A) KKT2 phosphorylates KKT8 of the KKT8 complex, while KKT3 phosphorylates KKT12 in vitro. In vitro kinase assay was performed using recombinant KKT8 complex with KKT2 or KKT3. The left panel shows phosphorylation detected by autoradiography, and the right panel shows the Coomassie Brilliant Blue staining. (B) KKT2 phosphorylates S381 of KKT8. In vitro kinase assay was performed on the KKT8 complex that has mutation in KKT8 S381A using full-length KKT2 or just the kinase domain. (C) Phosphorylation of KKT12 by KKT3 is abolished when T188 or S192 was mutated. (D) KKT8 S381 and the surrounding residues are well conserved among kinetoplastids. The putative phosphorylation motif of KKT2 is G-x-S*-Φ-x-F/Y-S-Φ, where S* gets phosphorylated and Φ represents hydrophobic residues. Note that the −2 position is Gly. (E) KKT12 T188, S192, and surrounding residues are well conserved among kinetoplastids. The putative phosphorylation motif of KKT3 is L-E/D-x-T*-A-G-E-S*-Φ-Φ-Φ-Φ, where either T* or S* (or both) gets phosphorylated. (F) KKT8^S381A^ mutant is functional. Growth curve of KKT8^S381A^-YFP with KKT8 3’UTR RNAi is shown. RNAi was induced with 1 µg/ml doxycycline to deplete the wild-type allele of KKT8. Control is an uninduced cell culture. Cell line: BAP1274

### Crystal structures of KKT2 DPB and KKT3 DPB reveal different surface charge distribution

We next characterized the C-terminal domain of KKT2 and KKT3, which has limited sequence similarities to polo boxes of PLK1 (Nerusheva and Akiyoshi, 2016). To gain insights into the function of these divergent polo boxes, we determined their high-resolution structures by X-ray crystallography. We obtained crystals of *T. congolense* KKT2^1030–1265^ (61.6% identical to *T. brucei* KKT2^1024–1260^) and determined its structure at 2.2 Å using molecular replacement with an AlphaFold2-predicted model of *T. congolense* KKT2 DPB as a search template, while the crystal structure of *T. brucei* KKT3 DPB was determined to 2.9 Å resolution using molecular replacement with a selenomethionine-derivatized crystal model (Figure 3A and Table 1). As expected from our previous sequence analysis (Nerusheva and Akiyoshi, 2016), structural homology searches using the DALI server (Holm, 2020) confirmed polo boxes of PLK1 as the closest structural homolog of KKT2/3 DPB (root mean square deviation [RMSD] between *T. congolense* KKT2 DPB and human PLK1: 6.3 Å across 109 Cα. RMSD between *T. brucei* KKT3 DPB vs human PLK1: 4.4 Å across 112 Cα) (Figure 3B and Table S2). However, some key residues involved in the phospho-peptide recognition are missing in KKT2/3 DPB as previously noted from their sequence analysis (Elia et al., 2003; Nerusheva and Akiyoshi, 2016) (Figure S4), suggesting either that KKT2/3 DPB is not a phosphorylation-dependent protein-protein interaction domain or that KKT2/3 DPB interacts with phosphorylated proteins in a distinct manner.

**Figure 3.**
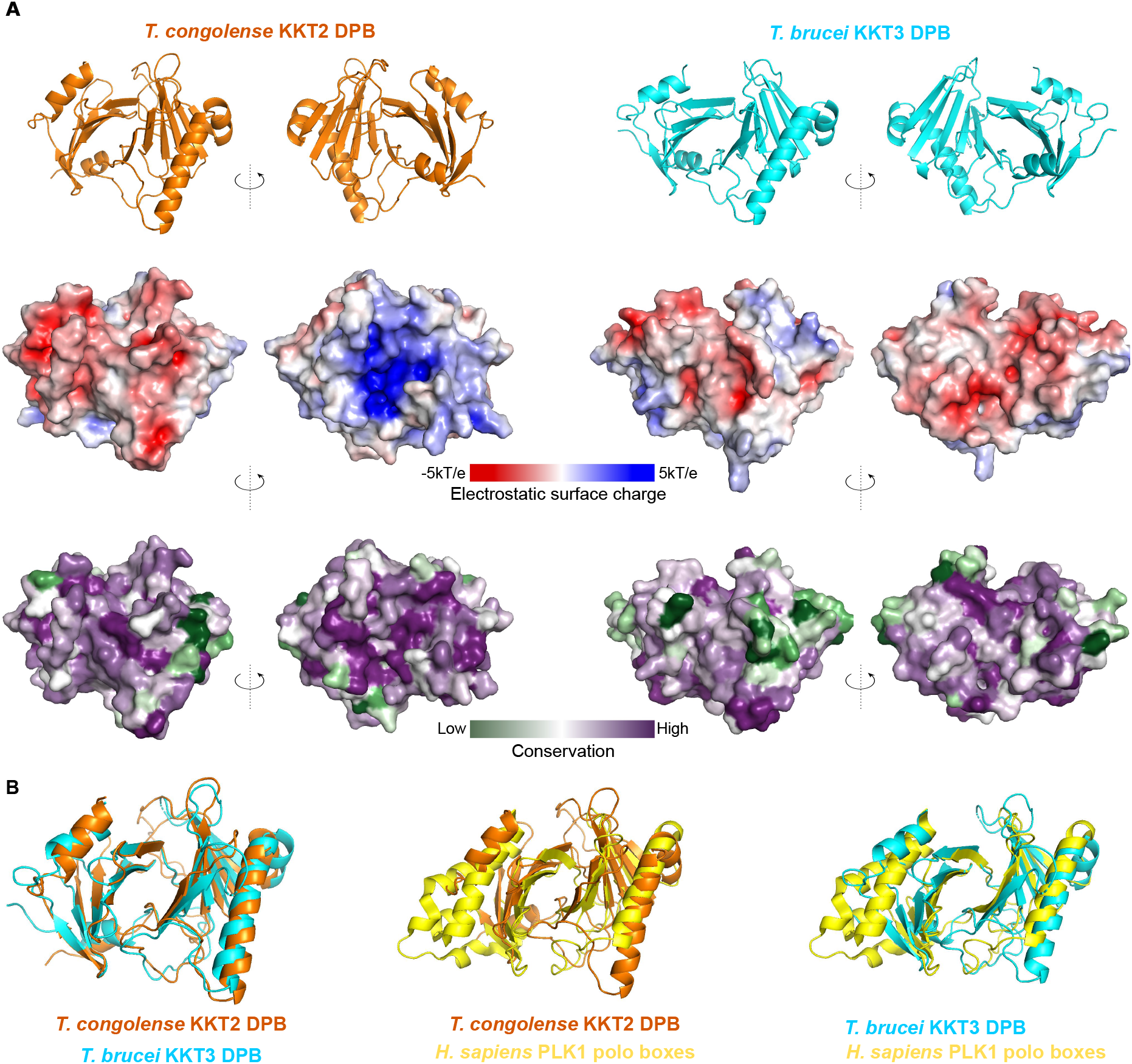
Crystal structures of KKT2 DPB and KKT3 DPB reveals differences. (A) Cartoon representation of the *T. congolense* KKT2 DPB (orange, left) and *T. brucei* KKT3 DPB (cyan, right), together with electrostatic surface potential and conservation of surface residues. Electrostatic surface potential of the DPBs was generated by the Adaptive Poisson– Boltzmann Solver (APBS) software (Jurrus et al., 2018). Conservation of surface residues was mapped based on sequence conservation using the ConSurf server (Ashkenazy et al., 2016). (B) Overlay of *T. congolense* KKT2 DPB and *T. brucei* KKT3 DPB (left), *T. congolense* KKT2 DPB and human PLK1 polo boxes (middle), and *T. brucei* KKT3 DPB and human PLK1 polo boxes (right). The structure of human PLK1 polo boxes is from (Qian et al., 2015) (PDB: 4X9R). These structures were aligned using “alignment” function in PyMOL.

**Table 1.** Data collection and refinement statistics*.

| Data collection | *T. congolense*<br>KKT2<sup>1030–1265</sup> | SeMet *T. brucei*<br>KKT3<sup>847–1058</sup> | Native *T. brucei*<br>KKT3<sup>847–1058</sup> |
| --- | --- | --- | --- |
| Beamline | Diamond Light Source I03 | Diamond Light Source I04 | Diamond Light Source I24 |
| Wavelength (Å) | 0.9763 | 0.9794 | 1.0072 |
| Space group (Z) | *P* 1 2<sub>1</sub> 1 | *P* 1 | *P* 1 |
| Unit cell | 48.70Å 57.13Å<br>83.67Å<br>90° 97.87° 90° | 45.06Å 101.14Å<br>102.53Å<br>93.39° 94.85°<br>98.15° | 45Å 52.49Å 102.15Å<br>84.11° 84.42° 73.25° |
| Resolution range (Å) | 47.04–2.20<br>(2.28–2.20) | 68.88–2.13<br>(2.17–2.13) | 50.68–2.92<br>(2.97–2.92) |
| Unique reflections | 23316 (2303) | 96388 (4664) | 19067 (926) |
| Completeness (%) | 95.80 (100) | 97.34 (94.59) | 99 (98.20) |
| Multiplicity | 6.8 (6.3) | 3.4 (3.4) | 3.4 (3.4) |
| *I*/σ*I* | 12.3 (0.7) | 5.6 (0.3) | 5.1 (1.4) |
| *R*<sub>meas</sub> | 0.098 (2.714) | 0.111 (1.279) | 0.183 (1.249) |
| CC<sub>1/2</sub> | 1.0 (0.4) | 0.97 (0.45) | 1.0 (0.54) |
| Wilson B-factor (Å<sup>2</sup>) | 32.68 | 40.29 | 63.19 |
| Refinement |  |  |  |
| No. reflections | 23299 (2300) |  | 18988 (383) |
| *R*<sub>work</sub> | 0.19 (0.28) |  | 0.25 (0.38) |
| *R*<sub>free</sub> | 0.24 (0.31) |  | 0.27 (0.42) |
| Number of atoms | 3674 |  | 6254 |
| Protein | 3489 |  | 6169 |
| Ligands | 0 |  | 34 |
| Solvent | 185 |  | 51 |
| RMS bonds (Å) | 0.007 |  | 0.008 |
| RMS angles (°) | 0.93 |  | 1.04 |
| Ramachandran favored (%) | 96.10 |  | 93.65 |
| Ramachandran allowed (%) | 3.90 |  | 5.72 |
| Ramachandran outliers (%) | 2.11 |  | 0.64 |
| Average B-factor (Å<sup>2</sup>) | 40.97 |  | 81.64 |

\*Statistics for the highest-resolution shell are shown in parentheses.

Although KKT2 DPB and KKT3 DPB have a highly similar backbone (RMSD: 2.2 Å across 129 Cα) (Figure 3B), they have distinct patterns of surface charge and conservation (Figure 3A, Figure S5A and S5B). One notable difference is that KKT2 DPB has positively charged patches (Figure 3A and Figure S5A). In addition, KKT2 DPB has a loop that is highly conserved among KKT2 homologs and is found even in early branching prokinetoplastids (Figure S5C and S5D). These differences raise a possibility that KKT2 DPB and KKT3 DPB have distinct functions.

### KKT3 DPB is essential for cell proliferation and can recruit KKT2 in trypanosomes

To examine functional importance of the divergent polo boxes of KKT2 and KKT3 for cell viability, we aimed to perform rescue experiments in trypanosomes using DPB mutants. Unfortunately, we could not analyze point mutants of KKT2/3 because they all had low protein level problems. Instead, we managed to obtain a deletion mutant that lacks KKT3 DPB (KKT3^ΔDPB^-YFP) and found that it failed to support cell growth when the remaining wild-type KKT3 protein was depleted by RNAi (Figure 4A). This result shows that KKT3 DPB is essential for the survival of procyclic trypanosome cells.

**Figure 4.**
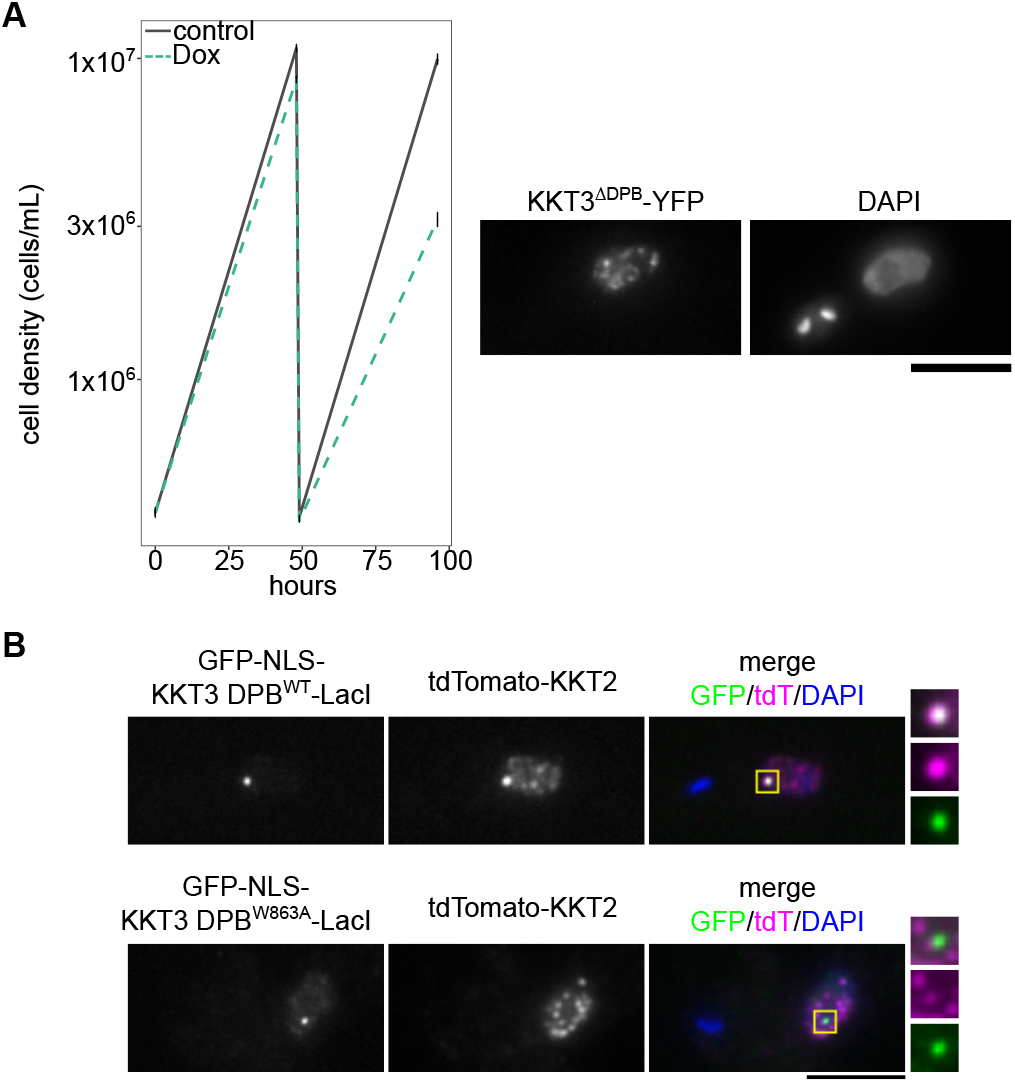
KKT3 DPB can recruit KKT2 to an ectopic locus. (A) KKT3^1–851^-YFP that lacks the DPB cannot support cell growth. Growth curve of KKT3^1–851^-YFP with KKT3 3’UTR RNAi is shown. RNAi was induced with 1 µg/ml doxycycline to deplete the untagged KKT3 allele. Control is an uninduced cell culture. Cell line: BAP2157 (B) KKT3 DPB, but not its W863A mutant, can recruit KKT2 to a non-centromeric locus in trypanosomes. Expression of GFP fusion proteins was induced in cells expressing tdTomato-KKT2 with 10 ng/ml doxycycline for 1 day. Cell lines: BAP2101, BAP2163. Bars = 5 µm.

We next aimed to reveal the function of KKT3 DPB. Our previous work showed that ectopically expressed KKT3 DPB co-purified with several kinetochore proteins including KKT2, KKT8, KKT7, and KKT1 (Marcianò et al., 2021). Using the LacO-LacI system (Ishii and Akiyoshi, 2020), we found that the GFP-KKT3 DPB-LacI fusion protein was able to recruit tdTomato-KKT2 to an ectopic locus (Figure 4B). This recruitment was abolished in the W863A mutant that is expected to affect the structural integrity of the KKT3 DPB protein (Figure 4B). These results show that KKT3 DPB is a protein-protein interaction domain that binds KKT2 either directly or indirectly.

### KKT2 DPB directly interacts with the C-terminal part of KKT1

Although KKT2 DPB has a sequence similarity with KKT3 DPB (25% identical), it remained unclear whether they have the same or distinct interaction partners. Indeed, our finding that the surface charges are quite different between these two proteins raised a possibility that they have distinct interaction partners. When expressed and immunoprecipitated from trypanosomes, KKT2 DPB co-purified with KKT1, KKT6, KKT7 and KKT8, which was abolished when KKT2 W1048 was mutated (Marcianò et al., 2021). Using the LacO-LacI tethering assay, we found that KKT2 DPB can recruit KKT1 to an ectopic locus in trypanosome cells, which was abolished in the W1048 mutant (Figure 5A). Based on AlphaFold2-based structure predictions, the N-terminal part of KKT1 (1–989) is predicted to contain putative HEAT repeats, while its C-terminal part (990–1594) is predicted to be largely disordered (Jumper et al., 2021; Wheeler, 2021; Mirdita et al., 2022; Varadi et al., 2022). While characterizing the KKT1 protein, we found that KKT2 was detected in the immunoprecipitates of KKT1C, but not KKT1N (Figure 5B and Table S3). Furthermore, our tethering assay showed that KKT1C was able to recruit KKT2 in trypanosome cells (Figure 5C). These results prompted us to test whether KKT2 DPB directly interacts with the C-terminal part of the KKT1 protein. Using recombinant proteins, we found that KKT2 DPB co-migrated with KKT1C^990–1594^ in analytical size-exclusion chromatography, which separates macromolecules based on their size and shape (Figure 5D). Interestingly, KKT1C eluted later in the presence of KKT2 DPB than KKT1C itself, suggesting that KKT1C, which is predicted to be largely disordered, may adopt a more compacted structure when it is bound to KKT2 DPB. Chemical crosslinking mass spectrometry assays identified a number of crosslinks between the two proteins (Figure 5E and Table S4). Taken together, these results establish that KKT2 DPB directly interacts with KKT1. In contrast, no apparent shift was observed for KKT3 DPB when mixed with KKT1C, suggesting that these proteins do not interact at least under the conditions used in our assay (Figure 5D).

**Figure 5.**
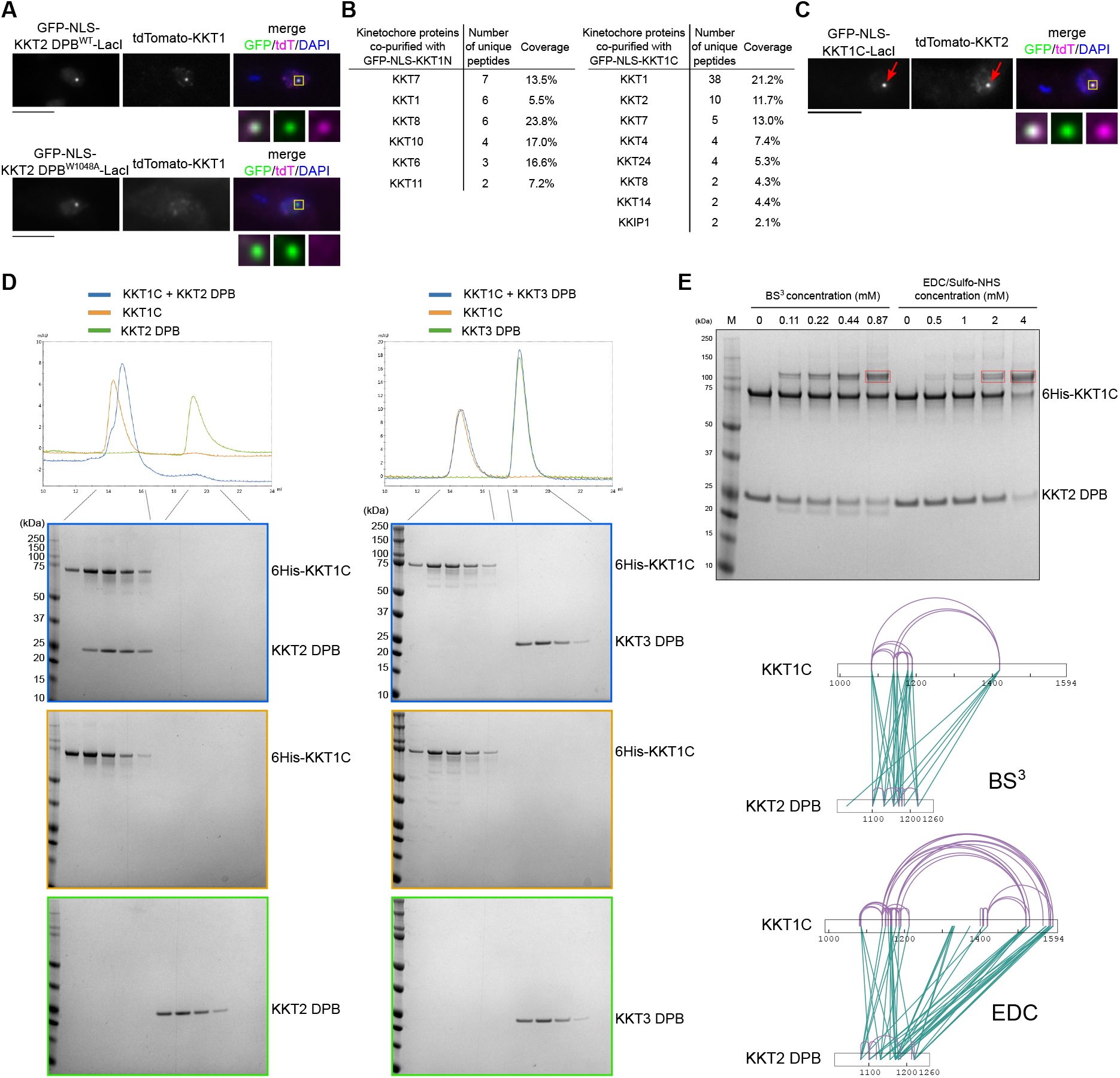
KKT2 DPB directly interacts with KKT1. (A) KKT2 DPB, but not its W1048A mutant, is sufficient to recruit KKT1 to a non-centromeric locus in trypanosomes. Cell lines: BAP1233, BAP2162 (B) Ectopically expressed GFP-NLS-KKT1C^990–1594^, but not GFP-NLS-KKT1N^2–989^, co-purifies with KKT2. See Table S3 for all proteins identified by mass spectrometry. Cell lines: BAP272, BAP273 (C) KKT1C^990–1594^ can recruit KKT2 to an ectopic locus in cells. Cell line: BAP2330. For A–C, expression of GFP fusion proteins was induced with 10 ng/ml doxycycline for 1 day. Bars = 5 µm. (D) KKT2 DPB^1024–1260^, not KKT3 DPB^832–1058^, forms a complex with KKT1C^990–1594^. Recombinant proteins were mixed on ice for 30 min, followed by analytical size exclusion chromatography on a Superose 6 10/300 column. (E) Crosslinking mass spectrometry of the KKT2 DPB/KKT1C complex using BS^3^ or EDC/Sulfo-NHS, showing extensive crosslinks between the two proteins. The bands highlighted in red boxes were digested with trypsin and analyzed by mass spectrometry. Green lines indicate inter-molecule crosslinks and purple lines indicate intra-molecule crosslinks. See Table S4 for all crosslinks identified by mass spectrometry.

### KKT1 forms a complex with KKT6

We next aimed to identify interaction partners for KKT1. In the immunoprecipitates of KKT1, several kinetochore proteins were identified (Akiyoshi and Gull, 2014). By co-expressing FLAG-tagged KKT1 with untagged KKT2, KKT5, KKT6, and KKT7, we found that KKT6 co-purified with FLAG-KKT1 (Figure S1C). To confirm this result, we next co-expressed only FLAG-KKT6 and KKT1 and found that these proteins still co-purified (Figure 6A). Crosslinking mass spectrometry identified many crosslinks between KKT1 and KKT6 (Figure 6B and Table S4). We therefore identified KKT6 as a direct interaction partner for KKT1.

**Figure 6.**
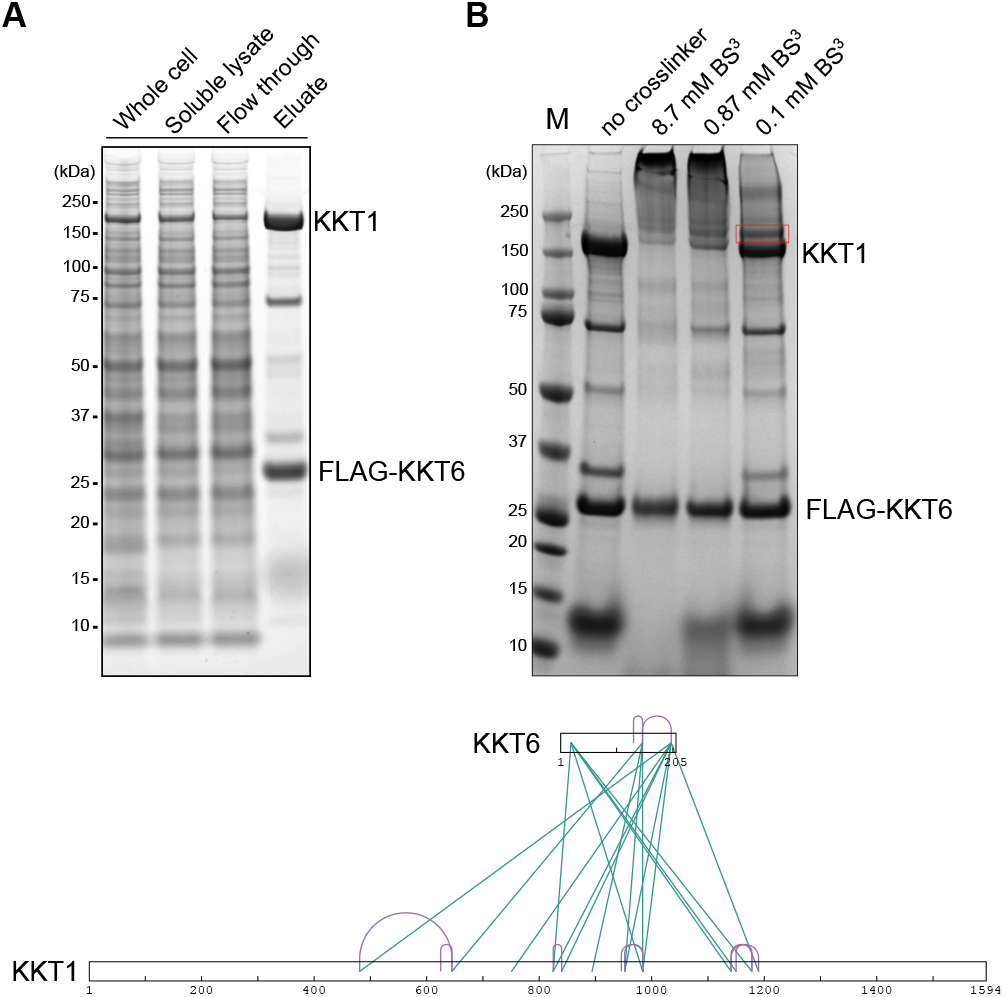
KKT1 interacts with KKT6. (A) KKT1 co-purifies with FLAG-KKT6. These proteins were co-expressed and purified from insect cells with anti-FLAG antibodies. (B) Crosslinking mass spectrometry of the KKT1/KKT6 complex using BS^3^. The band highlighted in the red box was digested with trypsin and analyzed by mass spectrometry. Green lines indicate inter-molecule crosslinks and purple lines indicate intra-molecule crosslinks. See Table S4 for all crosslinks identified by mass spectrometry.

## Discussion

Growing evidences point to the importance of KKT2 and KKT3 in the kinetoplastid kinetochores. These proteins have DNA-binding motifs, constitutively localize at kinetochores, and contribute to the recruitment of multiple kinetochore proteins, suggesting that KKT2 and KKT3 sit at the base of kinetoplastid kinetochores (Marcianò et al., 2021). These proteins share common ancestry with polo-like kinases and have an N-terminal protein kinase domain and C-terminal divergent polo boxes (Nerusheva and Akiyoshi, 2016). KKT2 and KKT3 additionally have a unique central domain that promotes centromere localization of these proteins (Marcianò et al., 2021). Homologs of KKT2/3 are found in essentially all sequenced kinetoplastids, although early branching prokinetoplastids have KKT2, rather than KKT3 (Butenko et al., 2020). We therefore speculated that ancestral kinetoplastids had only KKT2-like proteins that performed all necessary functions and that KKT3 in trypanosomatids represents a gene duplication product that became specialized in more efficient centromere localization (Marcianò et al., 2021). Our findings in this study highlight the difference between KKT2 and KKT3. The kinase activity of KKT2 is essential for accurate chromosome segregation, while that of KKT3 is dispensable for cell growth in the procyclic form *T. brucei*. Furthermore, KKT2 phosphorylates KKT8 of the KKT8 complex (which consists of KKT8/9/11/12), while KKT3 phosphorylates KKT12. Given that the KKT8 complex promotes kinetochore localization of the KKT10/19^CLK^ kinases (Ishii and Akiyoshi, 2020), these phosphorylation events might regulate the activity of KKT10/19.

Another difference between KKT2 and KKT3 proteins is their interaction partner. Our analysis identified KKT1 as the direct interaction partner for the DPB of KKT2, but not KKT3. However, it is important to mention that kinetochore localization of KKT1 was not significantly affected when KKT2 was depleted by RNAi but was affected when both KKT2 and KKT3 were depleted (Marcianò et al., 2021). These results imply redundancy in kinetochore recruitment of KKT1. It will be important to obtain a comprehensive interaction map of kinetoplastid kinetochore proteins. It will also be important to understand the molecular basis of the KKT2/KKT1 interaction in the future. It is worth mentioning that it represents the first direct protein-protein interaction identified between a constitutive kinetochore protein (KKT2) and a transient kinetochore protein that localizes from S phase onwards (KKT1). How is the recruitment of KKT1 onto KKT2 regulated during cell cycle? Although cyclin-dependent kinases play crucial roles in cell cycle-regulated recruitment of kinetochore proteins in other eukaryotes (Gascoigne and Cheeseman, 2013), our previous study found that kinetochore localization of KKT1 or other examined proteins were not affected by depletion of a mitotic cyclin in *T. brucei* (Hayashi and Akiyoshi, 2018). Based on the observation that the transcript of KKT1 is highly upregulated in S phase, it might be that KKT1 proteins are simply not available in G1 (Archer et al., 2011). Consistent with this possibility, the identified interaction between KKT2 DPB and KKT1C likely does not rely on phosphorylation because these proteins were expressed and purified from *E. coli*. Interestingly, not only KKT2/3 DPB but also polo boxes of the PLK1 homolog in *T. brucei* (called TbPLK that plays important roles in regulating cytoplasmic events such as the duplication of basal bodies and Golgi as well as cytokinesis) lack key residues involved in phospho-peptide binding in human PLK1 (Yu et al., 2012; McAllaster et al., 2015). Sequence analysis of PLK1 in *Naegleria* also suggests that its polo boxes unlikely act as a phospho-peptide binding domain (Nerusheva and Akiyoshi, 2016). We speculate that the last common ancestor of PLK1 might have been a phosphorylation-independent protein-protein interaction domain.

It has been shown that kinetoplastid kinetochores proteins are unique therapeutic targets against not only *T. brucei* but also *Leishmania* and *T. cruzi* (Nishino et al., 2013; Saldivia et al., 2020). Further understanding of molecular function and structure of the unique kinetoplastid kinetochore proteins could facilitate the development of specific and efficient drugs against neglected tropical diseases caused by kinetoplastid parasites.

## Materials and methods

### Primers, plasmids, bacmids, and synthetic DNA

All primers, plasmids, bacmids, and synthetic DNA used in this study are listed in Table S1. Their source or construction details are explained in Table S1. All constructs were sequence verified.

### Trypanosome cells

All trypanosome cell lines used in this study were derived from *T. brucei* SmOxP927 procyclic form cells (TREU 927/4 expressing T7 RNA polymerase and the tetracycline repressor to allow inducible expression) (Poon et al., 2012) and are listed in Table S1. Cells were grown at 28°C in SDM-79 medium supplemented with 10% (vol/vol) heat-inactivated fetal calf serum, 7.5 µg/ml hemin (Brun and Schönenberger, 1979), and appropriate drugs. Endogenous YFP tagging was performed using the pEnT5-Y vector (Kelly et al., 2007) or a PCR-based method (Dean et al., 2015). Endogenous tdTomato tagging was performed using pBA148 (Akiyoshi and Gull, 2014) and its derivatives. LacO-LacI tethering experiments were performed as described previously using the LacO array inserted at the rDNA locus (Landeira and Navarro, 2007; Ishii and Akiyoshi, 2020). Inducible expression of GFP-NLS fusion and GFP-NLS-LacI fusion proteins was carried out using pBA310 (Nerusheva and Akiyoshi, 2016) and pBA795 (Ishii and Akiyoshi, 2020), respectively. Cell growth was monitored using a CASY cell counter (Roche). Expression of GFP fusion proteins and RNAi were induced with doxycycline at a final concentration of 10 ng/mL and 1 µg/mL, respectively. One allele of KKT2 or KKT3 was deleted by a PCR-based method using a neomycin cassettes and primers listed in Table S1 (Merritt and Stuart, 2013; Ishii and Akiyoshi, 2020). To make a strain that lacks the KKT3 kinase domain, an N-terminal mNeonGreen tag was inserted between the kinase domain and the rest of the KKT3 coding sequence. Similarly, a C-terminal YFP tag was inserted just prior to the DPB of KKT3 to delete its DPB. Analog sensitive alleles or kinase dead mutant of KKT3 were made using endogenous tagging plasmids that contain appropriate mutations. The cell line that has kkt3Δ/KKT3-as1 (M109G) grew normally in the presence of 2.5 µM of 1NA-PP1, 1NM-PP1, or 3MB-PP1 (data not shown). To make deletion or mutant cell lines, transfected cells were selected with appropriate drugs and cloned by dispensing dilutions into 96-well plates and screened by PCR and/or DNA sequencing. All plasmids were linearized by *Not*I and transfected into trypanosomes by electroporation. Transfected cells were selected by the addition of 30 μg/mL G418 (Sigma), 25 μg/mL hygromycin (Sigma), 5 μg/mL phleomycin (Sigma), or 10 μg/mL blasticidin S (Insight biotechnology).

### Fluorescence microscopy

Cells were fixed with 4% paraformaldehyde for 5 min as described previously (Nerusheva and Akiyoshi, 2016), and images were captured at room temperature on a DeltaVision fluorescence microscope (Applied Precision) installed with softWoRx v.5.5 housed in the Oxford Micron facility essentially as described using a CoolSNAP HQ camera with 60x objective lenses (1.42 NA). Typically, ∼20 optical slices spaced 0.2 μm apart were collected. Images were processed in ImageJ/Fiji (Schneider et al., 2012). Maximum intensity projection images were generated by Fiji software (Schneider et al., 2012). Figures were made using Inkscape (The Inkscape Team).

### Multiple sequence alignment

Protein sequences and accession numbers for KKT2, KKT3, KKT8, KKT12 homologs used this study were retrieved from the TriTryp database (Aslett et al., 2010), UniProt (UniProt Consortium, 2019), or published studies (Tanifuji et al., 2017; Butenko et al., 2020; Tikhonenkov et al., 2021). Searches for homologous proteins were done using BLAST in the TriTryp database (Aslett et al., 2010). Searches for KKT2 and KKT3 homologs in Prokinetoplastina and Bodonida were done using hmmsearch on predicted proteome using manually prepared hmm profiles (HMMER version 3.0) (Eddy, 1998). Multiple sequence alignment was performed with MAFFT (L-INS-i method, version 7) (Katoh et al., 2019) and visualized with the Clustalx coloring scheme in Jalview (version 2.11) (Waterhouse et al., 2009).

### Expression and purification of *T. congolense* KKT2^1030–1265^

*T. congolense* KKT2^1030–1265^ used in this study was amplified from a synthesized gene fragment (BAG170) using BA3457 and BA3458 primers and cloned into the pRSFDuet-1 vector using NEBuilder Assembly 2x Master Mix (New England Biolabs) to make pBA2558 (*T. congolense* KKT2^1030–1265^ with an N-terminal tobacco etch virus [TEV]-cleavable hexahistidine [6His] tag). *E. coli* BL21(DE3) cells were transformed with ∼100 ng of plasmid DNA (pBA2558) and inoculated into 50 mL of 2xTY medium containing 50 µg/mL kanamycin and grown overnight at 37°C. In the next morning, each of the 6 L of 2xTY medium with 50 µg/mL of kanamycin was inoculated with 5 mL of the overnight culture and grown at 37°C with shaking (200 rpm) until the OD_600_ reached ∼0.8. Protein expression was induced with 0.2 mM IPTG for ∼16 hr at 20°C.

Cells were spun down at 3,400 g at 4°C and resuspended in 200 ml of lysis buffer (50 mM sodium phosphate, pH 7.5, 500 mM NaCl, and 10% glycerol) supplemented with protease inhibitors (20 μg/mL leupeptin, 20 μg/mL pepstatin, 20 μg/mL E-64, and 0.4 mM PMSF), benzonase nuclease (500 unit/L), and 0.5 mM TCEP. All subsequent steps were performed at 4°C. Bacterial cultures were mechanically disrupted using a French press (1 passage at 20,000 psi) and the soluble fraction was separated by centrifugation at 48,000 g for 30 min. Supernatants were loaded on 5 mL of TALON beads (Takara) pre-equilibrated with lysis buffer. Next, the beads were washed with ∼300 mL of the lysis buffer without protease inhibitors and proteins were eluted with 50 mM sodium phosphate pH 7.5, 500 mM NaCl, 10% glycerol, 250 mM imidazole and 0.5 mM TCEP. To cleave off the His-tag, samples were incubated with TEV protease in 1:50 w/w ratio overnight while being buffer-exchanged into 25 mM sodium phosphate, 250 mM NaCl, 5% glycerol, 5 mM imidazole, and 0.5 mM TCEP by dialysis. To increase the sample purity and remove the 6His tag, samples were re-loaded on TALON beads pre-equilibrated with dialysis buffer and the flow-through was collected. Next, the sample was concentrated using 10-kD MW Amicon concentrator (Millipore) and loaded on HiPrep Superdex 75 16/600 (GE Healthcare) columns to further purify and buffer exchange into 25 mM HEPES pH 7.5, 150 mM NaCl with 0.5 mM TCEP. Fractions containing KKT2 were pooled, concentrated to 12.6 mg/mL using a 10-kD MW Amicon concentrator (Millipore), and flash-frozen in liquid nitrogen for −80°C storage. Protein concentration was measured using absorbance at 280 nm and extinction coefficient calculated based on the protein sequence.

### Expression and purification of *T. brucei* KKT2 DPB, KKT3 DPB^832–1058^, KKT1C, and KKT2 kinase domain from *E. coli*

These recombinant proteins were purified based on the protocol used for *T. congolense* KKT2^1030–1265^ purification (see above) with the following modifications.

KKT2 DPB^1024–1260^ (pBA2240): Protein expression was induced at 30°C overnight using 24 L culture. After cell lysis with French press, Tween-20 was added to a final concentration of 0.2%. Dialysis was performed using buffer containing 25 mM HEPES, 200 mM NaCl, and 0.5 mM TCEP. Protein was then subjected to ion exchange chromatography using Resource S column with buffer A (25 mM HEPES pH 7.5, 0.5 mM TCEP) and buffer B (25 mM HEPES pH 7.5, 1 M NaCl, 0.5 mM TCEP), followed by size exclusion chromatography on HiPrep Superdex 75 16/600 (GE Healthcare) in 25 mM HEPES pH 7.5, 100 mM NaCl with 0.5 mM TCEP.

KKT3 DPB^832–1058^ (pBA2161): 6 L culture was used. Dialysis was performed using 50 mM sodium phosphate, pH 7.5, 500 mM NaCl, 10% glycerol, 5 mM imidazole, and 0.5 mM TCEP, followed by ion exchange chromatography using Resource Q column with buffer A (25 mM HEPES pH 7.5, 0.5 mM TCEP) and buffer B (25 mM HEPES pH 7.5, 1 M NaCl, 0.5 mM TCEP) and size exclusion chromatography on HiPrep Superdex 75 16/600 (GE Healthcare) in 25 mM HEPES pH 7.5, 150 mM NaCl with 0.5 mM TCEP.

6His-KKT1C^990–1594^ (pBA718): 6 L culture was used, protein expression was induced at 37°C for 4 hr, and the 6His tag was not cleaved for KKT1C. Immediately after elution from TALON beads, the protein was diluted with buffer A (25 mM HEPES pH 7.5, 0.5 mM TCEP) to a final concentration of 50 mM NaCl and further purified on ion exchange chromatography using Resource Q column with buffer A (25 mM HEPES pH 7.5, 0.5 mM TCEP) and buffer B (25 mM HEPES pH 7.5, 1 M NaCl, 0.5 mM TCEP). Size exclusion chromatography was done on HiPrep Superdex 200 16/600 (GE Healthcare) in 25 mM HEPES pH 7.5, 150 mM NaCl with 0.5 mM TCEP.

6His-KKT2 kinase domain (pBA318): 2.4 L culture of Rosetta (DE3) *E. coli* cells were grown at 16°C, and 1 ml of Talon beads were used. Eluted proteins (without TEV cleavage) were further purified on Superdex 200 16/600 (GE Healthcare) in 25 mM HEPES pH 7.5, 150 mM NaCl with 0.5 mM TCEP. Fractions that correspond to a monomer peak was collected and used for in vitro kinase assay.

### Expression and purification of *T. brucei* KKT3^847–1058^

*T. brucei* KKT3^847-1058^ was amplified from genomic DNA using primers BA676/BA600 and cloned into pNIC28-Bsa4 (Gileadi et al., 2008) using ligation-independent cloning to make pBA295 (*T. brucei* KKT3^847-1058^ with an N-terminal TEV-cleavable 6His tag). Recombinant protein was expressed in Rosetta (DE3) *E. coli* cells at 20°C using 12 L of 2xTY media. L-selenomethionine-labeled *T. brucei* KKT3^847-1058^ (SeMet *T. brucei* KKT3^847-1058^) recombinant protein was expressed in BL21(DE3) *E. coli* cells at 20°C using 6 L of medium base plus nutrient mix (SelenoMet medium, Molecular Dimensions). Cells were initially grown overnight in 2xTY media (100 ml), then centrifuged and resuspended twice with 60 ml of medium base plus nutrient mix. Each flask was inoculated with 10 ml of final resuspension, grown until OD_600_ of 0.6–0.8 and then supplemented with L-selenomethionine (65 mg per liter of media, Anagrade) and 0.2 mM IPTG.

Cells were harvested by centrifugation and resuspended in 25 mL lysis buffer (25 mM HEPES pH 7.5, 150 mM NaCl, 10 mM imidazole and 0.5 mM TCEP for unlabeled protein or 2 mM TCEP for labeled protein) per liter of culture. Cells were mechanically disrupted using French press (1 passage at 20,000 psi) and then centrifuged at 48,000 g for 30 min at 4°C. Tagged proteins were purified from lysate using 5 mL TALON beads (Takara), washed with 150 mL lysis buffer and eluted with 22 mL of elution buffer (25 mM HEPES pH 7.5, 150 mM NaCl, 250 mM imidazole and 0.5 mM TCEP for unlabeled protein or 2 mM TCEP for labeled protein), followed by overnight incubation with TEV protease in dialysis buffer (25 mM HEPES pH 7.5, 150 mM NaCl, 5 mM imidazole and 0.5 mM TCEP for unlabeled protein or 2 mM TCEP for labeled protein). Finally, 5 mL TALON beads column was used to remove the 6His tag and other contaminants from our sample. Cleaved protein was concentrated with an Amicon centrifugal filter with 10-kD cutoff (Merck) and then further purified by size exclusion chromatography using HiPrep Superdex 75 16/600 (GE Healthcare) pre-equilibrated with 25 mM HEPES (pH 7.5), 150 mM NaCl, and 0.5 mM TCEP for unlabeled protein or 4 mM TCEP for labeled protein. Fractions containing proteins were pooled together and concentrated to 12.2 mg/mL and 18.6 mg/mL for labeled protein and stored at −80°C. Protein concentration was measured by Bradford assay.

### Expression and purification of recombinant proteins from insect cells

3FLAG-KKT1 (bacmid pBA386), 3FLAG-KKT2 (bacmid pBA388), 3FLAG-KKT3 (bacmid pBA358), FLAG-KKT6/KKT1 (bacmid pBA828), 3FLAG-KKT1/KKT2 (bacmid pBA521), or 3FLAG-KKT1/KKT2/KKT5/KKT6/KKT7 (bacmid pBA523) were expressed in insect cells using the MultiBac baculovirus expression system (Bieniossek et al., 2012) (Geneva Biotech) using a protocol described previously (Llauró et al., 2018). Proteins were eluted in BH0.25 (25 mM HEPES, pH 7.5, 2 mM MgCl_2_, 0.1 mM EDTA, 0.5 mM EGTA, 10% glycerol, and 250 mM NaCl) supplemented with 0.5 mg/ml 3FLAG peptide (Sigma).

### Analytical size exclusion chromatography

8 µM of KKT2 DPB^1024–1260^ and 8 µM 6His-KKT1C were mixed for 30 min on ice. 7 µM of KKT3 DPB^832–1058^ and 7 µM 6His-KKT1C were mixed for 30 min on ice. All samples were in gel filtration buffer (25 mM HEPES pH 7.5, 150 mM NaCl with 0.5 mM TCEP). Analytical size exclusion chromatography was carried out on a Superose 6 10/300 (GE Healthcare) using gel filtration buffer on an ÄKTA pure system (GE Healthcare) at a flow rate of 0.5 ml/min at 4°C. Elution of proteins was monitored at 280 nm. 500 μL fractions were collected and analyzed by SDS-PAGE and Coomassie blue staining.

### Chemical crosslinking mass spectrometry

Crosslinking reactions were performed using BS^3^ and/or EDC/Sulfo-NHS essentially as described previously (Ludzia et al., 2021) using following samples: ∼2 µM of FLAG-KKT6/KKT1 in 25 mM HEPES pH 8.0, 2 mM MgCl_2_, 0.1 mM EDTA, 0.5 mM EGTA-KOH, 10% glycerol, 250 mM NaCl, 0.1% NP40, and 0.5 mg/ml 3FLAG peptide, or KKT2 DPB/KKT1C (taken from the analytical size exclusion chromatography experiment) in 25 mM HEPES pH 7.5, 150 mM NaCl with 0.5 mM TCEP. The crosslinked sample for the FLAG-KKT6/KKT1 complex was analyzed in the Advanced Proteomics Facility (https://www.proteomics.ox.ac.uk/). The gel band corresponding to crosslinked species was cut out, followed by in-gel trypsin digestion and LC-MS/MS analysis using a QExactive Orbitrap Mass Spectrometer (Thermo) as described previously (Ludzia et al., 2021). The crosslinked samples for KKT2 DPB/KKT1C complex were analyzed in the proteomics core facility at EMBL Heidelberg (https://www.embl.org/groups/proteomics/). The bands were subjected to in-gel digestion with trypsin (Savitski et al., 2014). Peptides were extracted from the gel pieces by sonication for 15 min, followed by centrifugation and supernatant collection. A solution of 50:50 water: acetonitrile, 1% formic acid (2x the volume of the gel pieces) was added for a second extraction and the samples were again sonicated for 15 minutes, centrifuged and the supernatant pooled with the first extract. The pooled supernatants were processed using speed vacuum centrifugation. The samples were dissolved in 10 µL of reconstitution buffer (96:4 water: acetonitrile, 1% formic acid) and analyzed by LC-MS/MS. An UltiMate 3000 RSLC nano LC system (Dionex) fitted with a trapping cartridge (µ-Precolumn C18 PepMap 100, 5µm, 300 µm i.d. x 5 mm, 100 Å) and an analytical column (nanoEase M/Z HSS T3 column 75 µm x 250 mm C18, 1.8 µm, 100 Å, Waters). Trapping was carried out with a constant flow of trapping solution (0.05% trifluoroacetic acid in water) at 30 µL/min onto the trapping column for 6 minutes. Subsequently, peptides were eluted via the analytical column running solvent A (0.1% formic acid in water) with a constant flow of 0.3 µL/min, with increasing percentage of solvent B (0.1% formic acid in acetonitrile). The outlet of the analytical column was coupled directly to an Orbitrap QExactive plus Mass Spectrometer (Thermo) using the Nanospray Flex ion source in positive ion mode. The peptides were introduced into the QExactive plus via a Pico-Tip Emitter 360 µm OD x 20 µm ID; 10 µm tip (New Objective) and an applied spray voltage of 2.2 kV. The capillary temperature was set at 275°C. Full mass scan was acquired with mass range 350-1500 m/z in profile mode with resolution of 70000. The filling time was set at maximum of 50 msec with a limitation of 3×10^6^ ions. Data dependent acquisition (DDA) was performed with the resolution of the Orbitrap set to 17500, with a fill time of 120 msec and a limitation of 5×10^4^ ions. A normalized collision energy of 30 was applied. Dynamic exclusion time of 30 sec was used. The peptide match algorithm was set to ‘preferred’ and charge exclusion ‘unassigned’, charge states 1 and 2 were excluded. MS^2^ data was acquired in centroid mode.

RAW MS files were searched by the pLink 2 software (Chen et al., 2019) using a FASTA database containing KKT1–20, KKT22–25, KKIP1, KKIP5, KKIP7, AUK1, CPC1, CPC2, KIN-A, KIN-B, and alpha/beta tubulins. Search parameters were as follows: maximum number of missed cleavages = 2, fixed modification = carbamidomethyl-Cys, variable modification Oxidation-Met. Precursor tolerance was set to 10 ppm. All the identified crosslinks are shown in Table S4 (FDR 5%). Crosslinks that have score < 1 x 10^-4^ were visualized in Figure 5 and Figure 6 using xiNET (Combe et al., 2015). All raw mass spectrometry files and custom database files used in this study have been deposited to the ProteomeXchange Consortium via the PRIDE partner repository (Perez-Riverol et al., 2019; Deutsch et al., 2020) with the dataset identifier PXD034039.

### Immunoprecipitation from trypanosomes and mass spectrometry

Expression of GFP-NLS-tagged KKT1N and KKT1C in trypanosomes was induced with 10 ng/mL doxycycline for 24 hours. Immunoprecipitation and mass spectrometry of these KKT1 fragments was performed with anti-GFP antibodies using a method we previously described (Ishii and Akiyoshi, 2020). Peptides were analyzed by electrospray tandem mass spectrometry over a 60-min gradient using QExactive (Thermo) at the Advanced Proteomics Facility (University of Oxford). RAW MS files were analyzed using MaxQuant version 2.0.1 (Cox and Mann, 2008) on a custom *T. brucei* proteome database that contains predicted proteins in TriTrypDB (TREU927, version 4) (Aslett et al., 2010) supplemented with predicted small proteins (Ericson et al., 2014; Parsons et al., 2015) with carbamidomethyl cysteine as a fixed modification and up to two missed cleavages allowed (protein FDR 1%). Default values were used except as follows. Oxidization (Met), phosphorylation (Ser, Thr, and Tyr), and acetylation (Lys) were searched as variable modifications. The first peptide tolerance was set to 10 ppm. Proteins identified with at least two peptides were considered as significant and shown in Table S3.

### In vitro kinase assay

10 µL of recombinant KKT8 complex (∼0.4 mg/mL in 50 mM sodium phosphate, pH 7.5, 500 mM NaCl, and 10% glycerol, and 250 mM imidazole) that was purified as described previously (Ishii and Akiyoshi, 2020) was mixed with 5 µL of recombinant kinase (3FLAG-KKT2: 0.22 mg/mL in 25 mM HEPES pH 8.0, 2 mM MgCl_2_, 0.1 mM EDTA, 0.5 mM EGTA-KOH, 10% glycerol, 250 mM NaCl, 0.1% NP40, and 0.5 mg/ml 3FLAG peptide. 3FLAG-KKT3: 0.5 mg/ml in 25 mM HEPES pH 8.0, 2 mM MgCl_2_, 0.1 mM EDTA, 0.5 mM EGTA-KOH, 10% glycerol, 250 mM NaCl, 0.1% NP40, and 0.5 mg/mL 3FLAG peptide. 6His-KKT2 kinase domain^60–382^: 0.4 mg/ml in 25 mM HEPES pH 7.5, 150 mM NaCl with 0.5 mM TCEP) into 1x kinase buffer (50 mM Tris-HCl pH 7.4, 1 mM DTT, 25 mM β-glycerophosphate, 5 mM MgCl_2_, 5 μCi [^32^P] ATP, and 10 μM ATP) in 25 µL reactions. The mixture was incubated at 30°C for 30 min, and the reaction was stopped by the addition of the LDS sample buffer (Thermo Fisher). The samples were run on an SDS-PAGE gel, which was stained with Coomassie Brilliant Blue R-250 (Bio-Rad) and subsequently dried and used for autoradiography using a phosphorimager screen. The signal was detected by an FLA 7000 scanner (GE Healthcare).

### Crystallization of *T. congolense* KKT2^1030–1265^ and *T. brucei* KKT3^847–1058^

All crystals were obtained in sitting drop vapor diffusion experiments in 96-well plates, using drops of 200 nL overall volume, mixing protein and mother liquor in a 1:1 ratio except for SeMet *T. brucei* KKT3^847-1058^ that was optimized in sitting drop vapor diffusion experiment in 48-well plate using drops of 400 nL overall volume, mixing protein and mother liquor in a 3:1 ratio. Crystals of *T. congolense* KKT2^1030–1265^ (10.7 mg/mL) were grown at 4°C in MIDAS HT-96 B5 solution (Molecular Dimensions) containing 0.1 M HEPES, pH 7.0, 8% w/v Polyvinyl alcohol, 10% v/v 1-Propanol. Crystals were briefly transferred into mother liquor prepared with addition of 25% glycerol prior to flash-cooling by plunging into liquid nitrogen. Crystals of native *T. brucei* KKT3^847-1058^ (12.2 mg/mL) were grown at 4°C in 25% PEG3350, 0.1 M Bis-Tris pH 5.5 and 0.1 M tri-sodium acetate pH 4.5. The crystals were briefly transferred into a cryoprotecting solution supplemented with 15% glycerol before flash cooling. Crystals of SeMet *T. brucei* KKT3^847-1058^ (18.6 mg/mL) were grown at 4°C in 27% PEG3350 and 0.05 M bis-tris pH 5.5. The crystals were briefly transferred into a cryoprotecting solution of 35% PEG3350 and 0.05 M Bis-Tris pH 5.5 before flash cooling.

### Diffraction data collection and structure determination

X-ray diffraction data from *T. congolense* KKT2^1030–1265^ were collected at the i03 beamline at the Diamond Light Source (Harwell, UK). The structure was solved using PHASER (McCoy, 2017), a molecular replacement software, with AlphaFold2-predicted structure of *T. congolense* KKT2^1030–1265^ as a model. Following the molecular replacement, the initial model building was done with BUCCANEER (Cowtan, 2006). Further manual model building and refinement were completed iteratively using COOT (Emsley et al., 2010) and PHENIX (Liebschner et al., 2019). Prior to the final refinement, the data were scaled to 2.2 Å resolution.

SeMet *T. brucei* KKT3^847-1058^ X-ray diffraction data were collected at the I04 beam line at Diamond Light Source at the selenium K-edge wavelength (0.9795 Å) and processed using the Xia2 pipeline (Winter, 2010), with DIALS for indexing and integration (Winter et al., 2018) and AIMLESS for scaling to 2.13 Å (Evans and Murshudov, 2013). Initial phases and model were obtained with the Big EP pipeline (Sikharulidze et al., 2016) using autoSHARP (Vonrhein et al., 2007), Phenix AutoSol (Terwilliger et al., 2009) and Crank2 (Skubák and Pannu, 2013). The structure was then completed by using BUCCANEER (Cowtan, 2006) followed by alternate cycles of model building in Coot and refinement in autoBUSTER (Blanc et al., 2004; Bricogne et al., 2017). Native *T. brucei* KKT3^847-1058^ X-ray diffraction data were collected at the I24 beam line at Diamond Light Source and processed using Xia2 pipeline, with DIALS for indexing and integration and AIMLESS for scaling to 2.92 Å. Initial phases have been determined by molecular replacement with PHASER using a SeMet *T. brucei* KKT3^847-1058^ structure as search model. The structure was completed with alternate cycles of model building in Coot and refinement in autoBUSTER. All images were made with PyMOL (version 2.5, Schrödinger). Protein coordinates have been deposited in the RCSB protein data bank with accession numbers 8A0J (*T. congolense* KKT2 DPB) and 8A0K (*T. brucei* KKT3 DPB).

## Acknowledgments

We thank the crystallography facility manager Edward Lowe, Micron Advanced Bioimaging Unit, Advanced Proteomics Facility in Oxford, and the proteomics core facility at EMBL, especially Mandy Rettel and Jennifer Schwarz, for their support. We also thank Miguel Navarro (Instituto de Parasitologıá y Biomedicina López-Neyra, Consejo Superior de Investigaciones Cientificas, Spain) for providing the pMig75 plasmid, Song Tan (Department of Biochemistry & Molecular Biology, The Pennsylvania State University, USA) for the pST44 plasmid, and Sam Dean (Warwick Medical School, University of Warwick, UK) for pPOT plasmids. M. Ishii was supported by a long-term fellowship from the TOYOBO Biotechnology Foundation. P. Ludzia was supported by the Boehringer Ingelheim Fonds. B. Akiyoshi was supported by a Wellcome Trust Senior Research Fellowship (grant 210622/Z/18/Z) and the European Molecular Biology Organization Young Investigator Program. The authors declare no competing financial interests.

## Rights retention

This research was funded in whole or in part by Wellcome Trust (grant 210622/Z/18/Z). For the purpose of Open Access, the author has applied a CC BY public copyright licence to any Author Accepted Manuscript (AAM) version arising from this submission.

## Author contributions

M. Ishii performed LacO-LacI tethering, RNAi, and rescue experiments in trypanosomes. P. Ludzia purified various recombinant proteins, identified interaction between KKT1C and KKT2 DPB, and performed crosslinking mass spectrometry. P. Ludzia and W. Allen solved the KKT2 DPB crystal structure. G. Marcianò solved the KKT3 DPB crystal structure. O.O. Nerusheva made analog-sensitive KKT2 and KKT3 alleles as well as KKT3 mutants. B. Akiyoshi performed in vitro kinase assays and immunoprecipitation of KKT1 fragments from trypanosomes. B. Akiyoshi wrote the manuscript. M. Ishii, P. Ludzia, and G. Marcianò edited the manuscript.

## Supplementary material

### Supplementary tables (Excel files)

Table S1. List of trypanosome cell lines, plasmids, bacmids, primers, and synthetic DNA used in this study

Table S2. DALI search results for *T. congolense* KKT2 DPB and *T. brucei* KKT3 DPB

Table S3. List of proteins identified in the immunoprecipitates of GFP-KKT1N and GFP-KKT1C by mass spectrometry and MaxQuant analysis

Table S4. List of crosslinks for the KKT2 DPB/KKT1C complex (BS^3^ and EDC) and KKT1/KKT6 complex (BS^3^) identified by mass spectrometry and pLink2 analysis

### Supplemental figures

**Figure S1.**
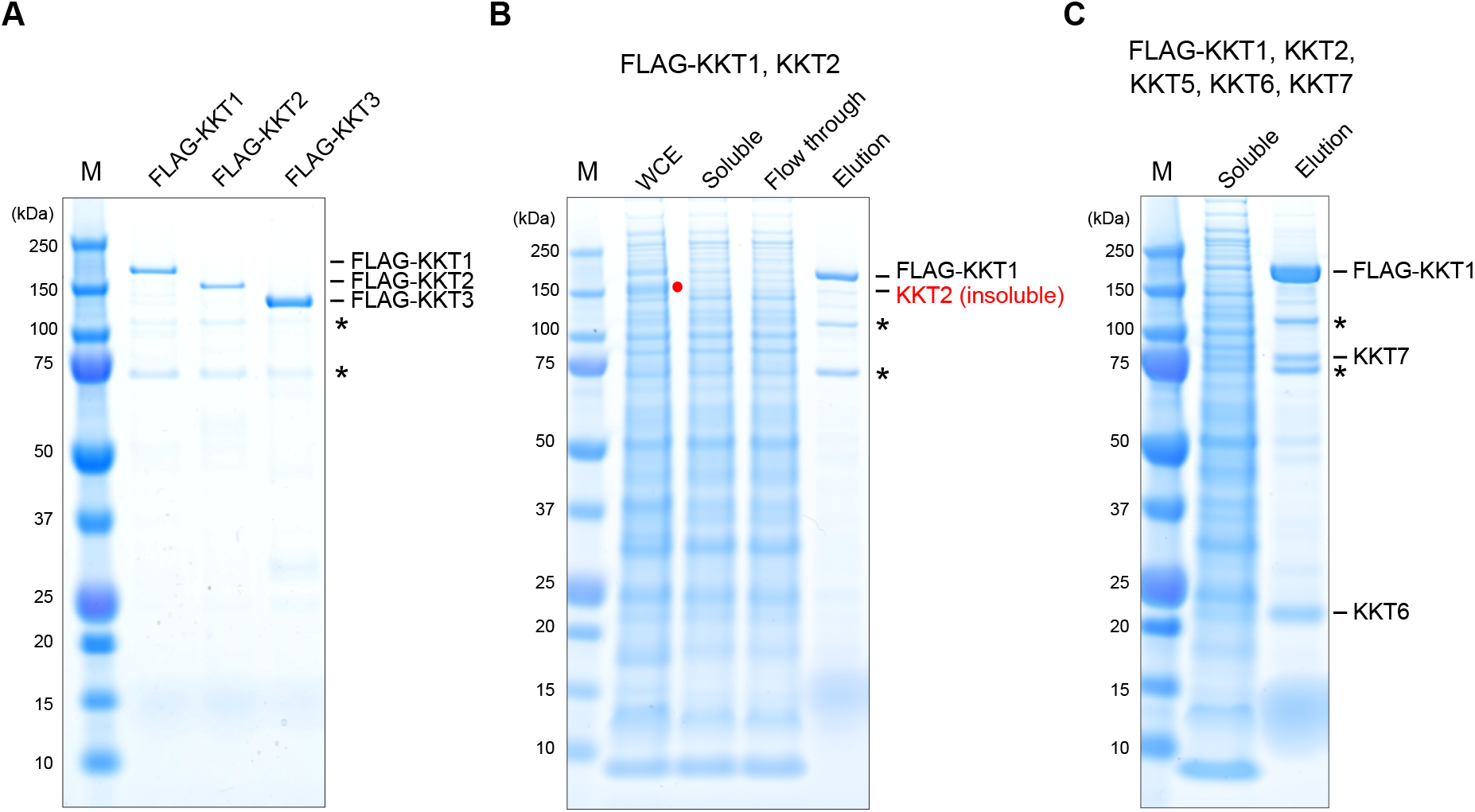
Purification of recombinant kinetochore proteins from insect cells. Indicated proteins were expressed and purified from insect cells using the MultiBac baculovirus expression system. FLAG-tagged proteins were purified using anti-FLAG antibodies and eluted from beads with FLAG peptides. Coomassie-stained SDS-PAGE gels are shown. Asterisks indicate common contaminants. (A) Purification of FLAG-KKT1, FLAG-KKT2, and FLAG-KKT3. (B) Purification of FLAG-KKT1 from insect cells that also express KKT2. Note that untagged KKT2 was largely insoluble (shown in red). WCE stands for whole cell extract. (C) Purification of FLAG-KKT1 from insect cells that also express KKT2, KKT5, KKT6, and KKT7. A near-stoichiometric amount of KKT6 co-purified with FLAG-KKT1, while KKT7 was sub-stoichiometric. Identity of KKT6 and KKT7 was confirmed by mass spectrometry of indicated bands.

**Figure S2.**
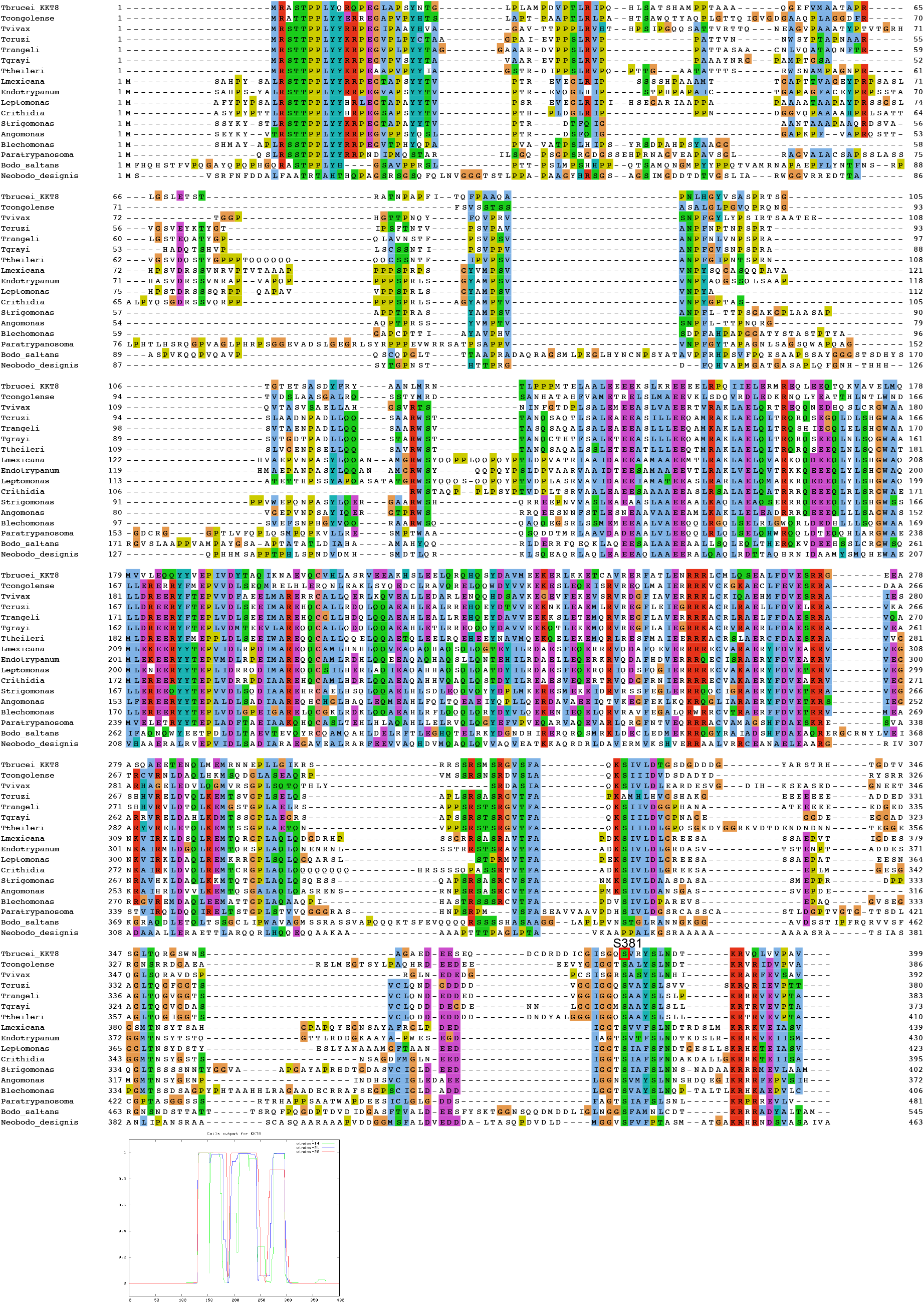
Multiple sequence alignment of KKT8 and its coiled-coil prediction. *Tb*KKT8 S381, the major target of KKT2, is highly conserved among kinetoplastids. Coiled-coil prediction was performed using coils server (Lupas et al., 1991).

**Figure S3.**
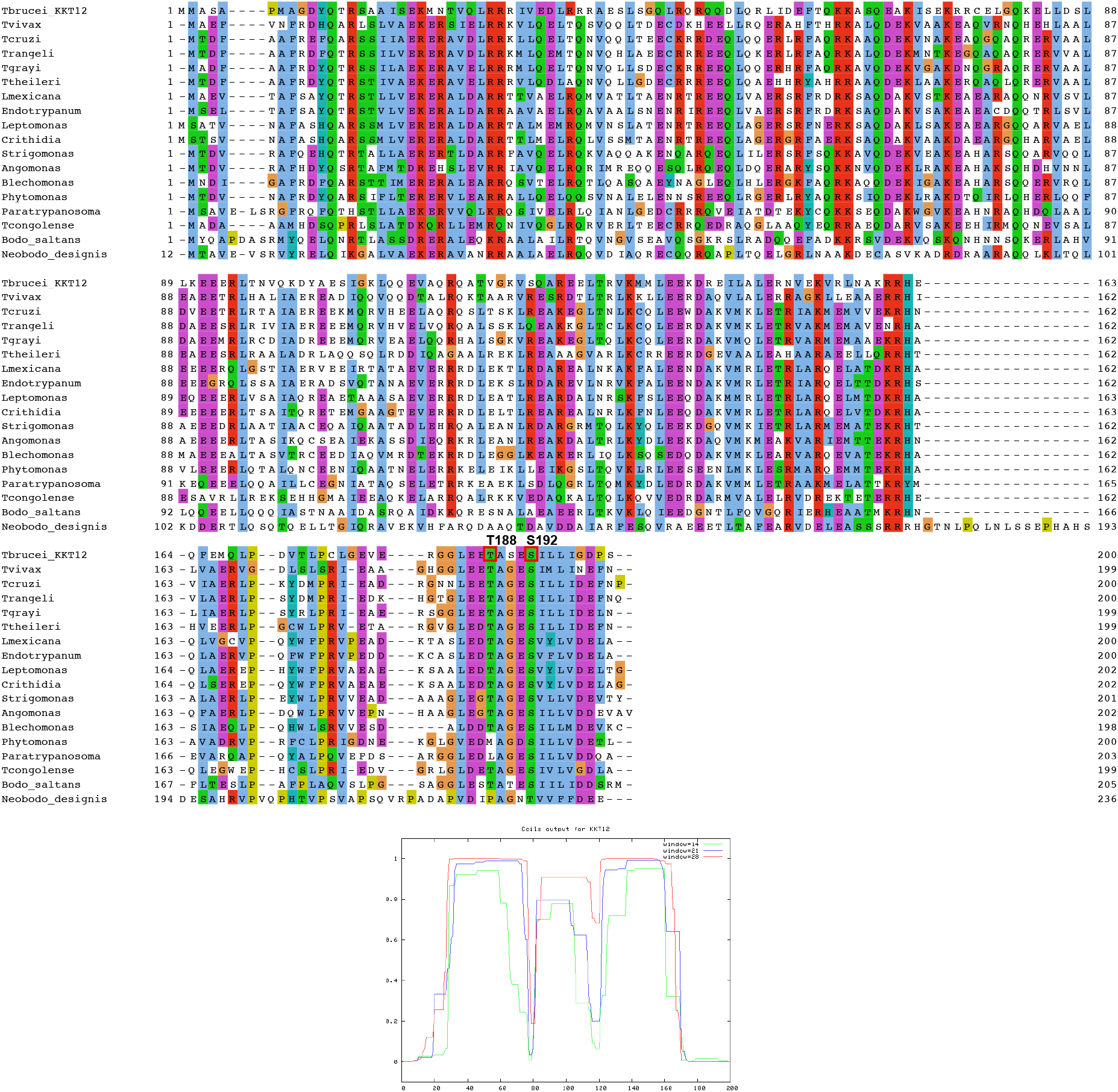
Multiple sequence alignment of KKT12 and its coiled-coil prediction. Residues T188 and S192 in *Tb*KKT12 are highlighted.

**Figure S4.**
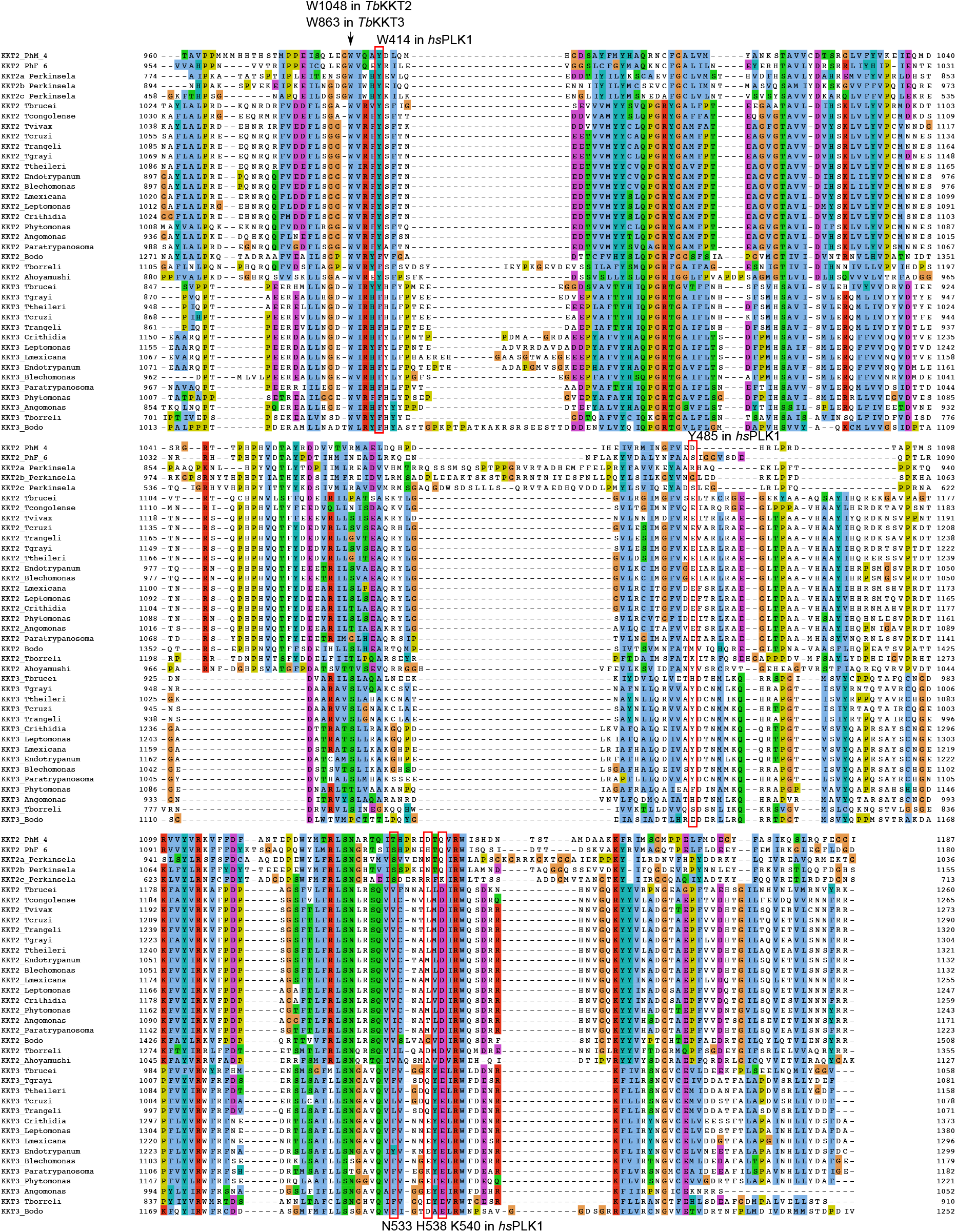
Multiple sequence alignment of KKT2 and KKT3. Highly conserved residues *Tb*KKT2 W1048 and *Tb*KKT3 W863 mutated in this study are indicated by arrow. Red boxes indicate equivalent positions of human PLK1 residues that play important roles in recognition of phospho-peptides (Elia et al., 2003) but are not conserved in KKT2 or KKT3. PhM_4 (*Papus ankaliazontas*), PhF_6 (*Apiculatamorpha spiralis*), and *Perkinsela* belong to Prokinetoplastina, the most divergent group of kinetoplastids (Tanifuji et al., 2017; Butenko et al., 2020; Tikhonenkov et al., 2021)

**Figure S5.**
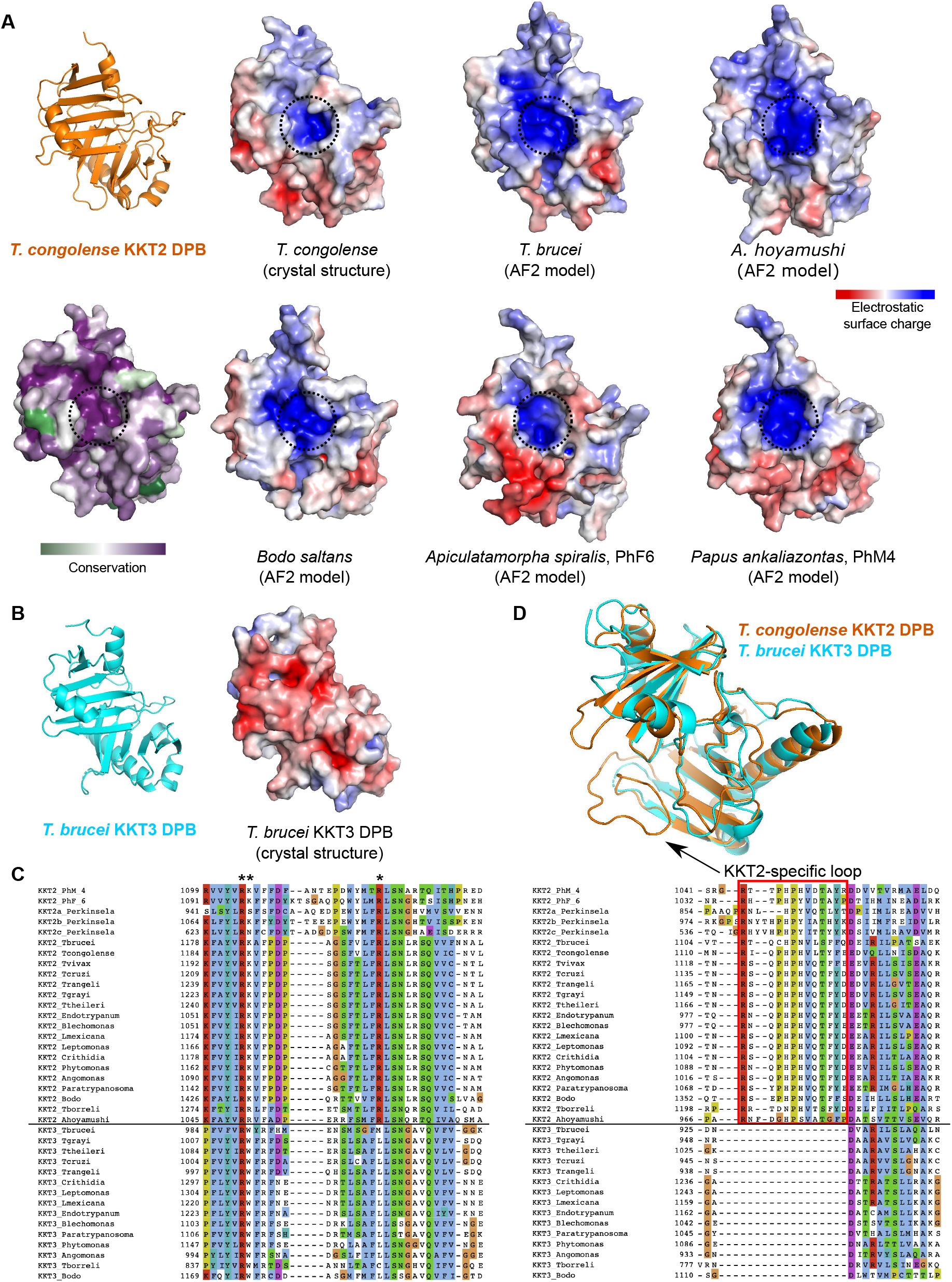
KKT2 has a highly conserved basic patch and a unique loop insertion. (A) Electrostatic surface potential of the *T. congolense* KKT2 DPB crystal structure or AlphaFold2-models from various kinetoplastids. Electrostatic surface potential was calculated by the APBS software and is displayed in the range of −5 to +5 kT/e (Jurrus et al., 2018). The basic surface patch that is conserved among kinetoplastids is highlighted by dotted circle. (B) Electrostatic surface potential of the *T. brucei* KKT3 DPB crystal structure, showing difference in surface charge potential. (C) Multiple sequence alignment of KKT2 and KKT3. Positively charged residues that contribute to the basic patch of KKT2 in (A) are highlighted by asterisks. Note that R1197 in *Tb*KKT2 is unique to KKT2 proteins and is strictly conserved. (D) Top: Overlay and of *T. congolense* KKT2 DPB (orange) and *T. brucei* KKT3 DPB (cyan) reveals KKT2-specific loops. Bottom: Multiple sequence alignment of KKT2 and KKT3 shows that the highlighted loop is highly conserved among KKT2 proteins.

